# Male-specific analgesic effects of minocycline in sickle cell disease are mediated by microglia and the microbiome

**DOI:** 10.1101/2025.08.26.672427

**Authors:** Juanna M. John, Zulmary Manjarres, Nurfatihah I. Zulkifly, Ashley N. Plumb, McKenna L. Pratt, Katelyn E. Sadler

## Abstract

Over 50% of individuals with sickle cell disease (SCD) experience chronic pain that is phenotypically distinct from their acute, vaso-occlusive crisis pain. Chronic SCD pain is commonly managed with opioid-based drugs that are associated with unwanted side effects, incomplete pain relief, and – in this population – accessibility issues. Thus, new treatments for chronic SCD pain are desperately needed. Here, we examined the analgesic efficacy of acute minocycline treatment in transgenic SCD mice. SCD mice exhibit gut dysbiosis and chronic inflammation. Therefore, we hypothesized that minocycline would provide robust analgesia in this model given the drug’s antibiotic and anti-inflammatory properties respectively. Six days of minocycline treatment reversed chronic mechanical hypersensitivity only in male SCD mice. We identified two potential mechanisms underlying these sex-specific effects. First, we observed increased microgliosis only in the dorsal horn of male SCD mice. Minocycline treatment had opposite effects on microglial number in male and female SCD spinal cords. Second, minocycline treatment altered the gut microbiota in a sex-specific fashion; fecal microbiota transplant (FMT) from minocycline-treated female SCD mice induced widespread pain in recipients whereas FMT from minocycline-treated male SCD mice did not. In summary, these experiments highlight novel sex-specific mechanisms of minocycline analgesia and support future exploration of minocycline use for SCD pain management, but only in male patients.

## 1. Introduction

Chronic pain is a debilitating symptom experienced by individuals with sickle cell disease (SCD) [11]. Worldwide, approximately 7.74 million individuals suffer from this genetic blood disorder [64] which – in addition to pain – is characterized by hemolytic anemia, chronic inflammation [15], and changes in the gut microbiome [8,9,18,20,38,44,57]. Individuals with SCD describe their chronic pain as phenotypically distinct from the pain they experience during an acute vaso-occlusive episode [17,24]. Verbal descriptions of chronic SCD pain include words that suggest both nociceptive (*e*.*g*., throbbing, piercing, crushing) and neuropathic (*e*.*g*., aching, shooting, stabbing) underlying mechanisms [6,28]. As such, the most successful analgesic strategies for alleviating chronic SCD pain are likely to be those that target multiple pathological processes. Current guidelines for pharmacological management of chronic SCD pain include use of non-steroidal anti-inflammatory drugs (NSAID), serotonin and norepinephrine reuptake inhibitors, tricyclic antidepressants, and gabapentinoids [5]. Unfortunately, these drugs and opioid-based therapies, which have additional unwanted side effects, often fail to completely alleviate chronic SCD pain. Thus, additional drug classes with multiple modes of analgesic action should be considered for use in individuals with SCD.

Chronic inflammation and gut dysbiosis are two hallmarks of SCD that contribute to chronic pain and likely develop as a result of disease-associated hemolysis and vaso-occlusion [15]. Upon sickling, erythrocytes in both patients and mice with SCD release pro-inflammatory compounds (*e*.*g*., reactive oxygen species), adhere to the vascular endothelium and circulating immune cells, and ultimately, undergo hemolysis which results in the elevation of circulating damage associated molecular patterns (DAMPs; *e*.*g*., heme, ATP). These red cell-derived DAMPs can activate microglia – resident immune cells of the CNS that are increased in both number and activation status in the SCD mouse spinal cord dorsal horn [65]. In addition, work from our own lab recently demonstrated that hemolysis-associated metabolites in the gastrointestinal tract alter the gut microbiota and drive chronic pain in SCD mice [57]. Until curative SCD therapies become widely available and erythrocyte sickling can be prevented from the time of birth, analgesic therapies that simultaneously alter the gut microbiome and limit inflammation may provide significant relief for those suffering from SCD pain.

To this end, we examined the analgesic efficacy of minocycline in SCD mice. Minocycline is an inexpensive, widely available drug that was originally characterized as a broad spectrum tetracycline antibiotic [49,55]. Tetracyclines limit bacterial growth by binding to the 30S bacterial ribosome subunit, thus inhibiting protein translation. Minocycline activity is not limited to prokaryotic cells, however; minocycline modulation of host immune cell function also limits inflammation in a variety of clinical conditions and rodent models. In these studies, we administered minocycline to male and female SCD mice for six days then assessed pain-like behaviors, changes in the gut microbiota composition, and microglial activation in the spinal cord dorsal horn – a key relay in CNS nociception networks. Our goal for this study was not only to assess the analgesic efficacy of minocycline in SCD, but to further understand the cellular basis – both prokaryotic and eukaryotic – of chronic pain in SCD.

## 2. Methods

### 2.1 Mice

These experiments primarily used the Townes mouse model of sickle cell disease (SCD) [71]. In this model, endogenous murine α- and β-globin genes are knocked out and replaced with human γ-, α- and either normal (‘A’) or sickle (‘S’) β-globin genes. Mice homozygous for sickle β-globin (HbSS) display many SCD characteristics including sickled erythrocytes, hemolytic anemia [71], hypoxia-reoxygenation (H/R)-inducible acute pain [10], spontaneous chronic pain [50,58], and persistent hypersensitivity to mechanical and cold stimuli [33,40,61,77]. Mice homozygous for wildtype β-globin (HbAA) are used as controls. Male and female HbAA and HbSS mice aged 8-20 weeks were used in experiments. C57BL/6J male and female mice, aged 8-12 weeks, were used as fecal material transplant recipients. All mice were bred in-house and had *ad libitum* access to food and water throughout the duration of all experiments. Mice were randomized to treatment groups. All animal experiments and procedures in this study were done following approval from the Institutional Animal Care and Use Committee of the University of Texas at Dallas (protocol #2022-0088).

### 2.2 Pain-like behavioral assays

Experimenters were blinded to mouse genotype and/or treatment group during behavioral testing and analysis. All testing was completed in Plexiglass behavior chambers (10 x 10 x 20 cm); chambers were placed on a raised wire mesh grid for mechanical assays and a ¼’ thick piece of glass for dry ice testing. Animals were habituated to the testing environment for at least one hour before testing. The experimenter was present for at least the last 30 min of this habituation period.

#### 2.2.1 Von Frey mechanical hypersensitivity testing

The up-down von Frey assay was used to measure punctate mechanical hypersensitivity as previously described [13]. Hindpaw 50% withdrawal thresholds were recorded by applying calibrated von Frey filaments (0.02 – 2.56g) at a 30-degree angle to the center of the plantar surface of each hindpaw for 2 seconds. If the mouse raised its stimulated paw from the mesh, it was classified as a response. If the first filament elicited a withdrawal, the next lower force filament was then used to stimulate the same paw. Conversely, if a withdrawal was not detected with the first filament, the next larger force filament was used. After the first series of opposite reactions (*i.e*., withdrawal response followed by no response or vice versa), each hindpaw was stimulated four more times. The 50% withdrawal threshold for each hindpaw was separately calculated as described [21] before being averaged for each animal.

#### 2.2.2 Needle mechanical hyperalgesia testing

Hindpaw needle stimulation was used to measure mechanical hyperalgesia as previously outlined [34]. Needle responses were elicited by poking the center of the plantar surface of each hindpaw with a 27Ga spinal needle. Subsequent responses were classified into three categories: null, normal, and nocifensive. Null responses were those in which the animal did not respond to stimulation. Normal responses were those in which the animal first lifted then returned its paw to the wire mesh. Responses were classified as nocifensive when the animal raised its paw and additionally shook, licked, or persistently elevated the paw. Needle testing was repeated 5 times on each paw and then response frequency and types were summed between paws.

#### 2.2.3 Brush dynamic mechanical allodynia testing

Hindpaw brush stimulation was used to measured dynamic mechanical allodynia as previously described [16]. Brush responses were elicited by lightly dragging a 10/0 short liner paintbrush across the entire glabrous surface of each hindpaw from heel to toe. Responses to dynamic brush stimulation were characterized and analyzed like the needle assessment.

#### 2.2.4 Plantar dry ice cold hypersensitivity testing

The plantar dry ice assay measured hypersensitivity to cold stimulation as previously described [7]. Dry ice was crushed into a powder, loaded into a plastic applicator, then applied to the glass directly below each hindpaw; the time taken to withdraw each paw (withdrawal latency; not to exceed 20s) was recorded. The withdrawal latency was measured 5 times for each paw then averaged between paws to calculate the overall withdrawal latency for a single mouse.

### 2.3 Animal treatments

#### 2.3.1 Minocycline administration

Townes HbAA and HbSS mice had *ad libitum* access to 100 mg/kg minocycline treatment (Santa Cruz Biotechnology, Cat# sc-203339B) for 6 days; similar doses of minocycline have previously been shown to decrease immune cell infiltration in the HbSS central nervous system [19]. Water bottles were weighed and refilled every other day. Pain-like behavior tests were performed before and after 6 days of vehicle (water) or minocycline treatment.

#### 2.3.2 Fecal microbiota transplant (FMT) paradigm

Fecal material was collected immediately upon defecation from vehicle or minocycline-treated (days 4-10 of treatment) HbAA and HbSS mice. Fecal donors were housed in 2 separate cages to limit cage effects frequently observed in microbiome studies. Upon defecation, feces were frozen at -80°C or immediately homogenized in vivarium drinking water (1 pellet /1mL water). Homogenized fecal material was administered to naïve, sex-matched C57BL/6 mice every other day for 7 days (4 FMTs total; 200µL p.o.). Pain-like behavior assessments were performed prior to FMT (B: baseline), on the day between the 3^rd^ and 4^th^ FMTs (FMT), 7 and 14 days following the final FMT.

### 2.4 Gross anatomical assessments

One day following the completion of behavior testing, animals were euthanized via sodium pentobarbital injection. Tissues/samples of interest (spleen, small intestine, colon, cecum, fecal material, and spinal cord) were collected and processed as described below. The spleen and cecum were removed and immediately weighed. The small intestine and colon were removed, cleaned of residual mesentery, and measured for length. Fecal samples and spinal cord collection methods are outlined below.

### 2.5 Spinal cord Iba1+ immunofluorescent staining

#### 2.5.1 Spinal cord tissue processing

After mice were euthanized, spinal cords were dissected and drop fixed in 4% PFA overnight at 4°C. Spinal cords were cryopreserved in 30% sucrose at 4°C for 48hr then embedded in optimal cutting temperature (OCT) media before being sectioned on a cryostat. Tissue was sliced transversally at 30µm and stored in 1X PBS (4°C) before staining.

#### 2.5.2 Iba1 immunofluorescent staining

Sections (6 per animal) containing lumbar spinal cord (L4-L6) were mounted onto SuperFrost Plus slides then incubated in blocking solution (3% normal goat serum, 2% bovine serum albumin, 1% Triton-X100, 0.05% Tween-20, and 1X PBS) for 1hr. Spinal cord sections were immunostained with a rabbit anti-Iba1 (1:500; Wako Fujifilm, Cat# 019-19741). After washing several times, sections were incubated with a AlexaFluor-488 goat anti rabbit IgG (H+L) (1:500, Invitrogen, Cat#: A11008) and Hoechst 33342 (1:1000, Invitrogen, Cat# H3570,). All sections were mounted with ProLong Gold antifade reagent.

#### 2.5.3 Confocal imaging and analysis of microglia

For analysis of Iba1, three images of either left or right dorsal horn were collected at 20X per animal using a FV-4000 (Olympus; UPlanXApo 20X/0.80/0.17/OFN26.5) confocal microscope. Note, no differences in Iba1 staining were observed between left and right dorsal horn within a given animal. Images were digitized with confocal Z-stacks and maximum intensity projection was analyzed. The number of Iba1+ cells in laminae I, II, and III were counted manually using the cell counter plugin in Image J (http://imagej.nih.gov). The investigator was blinded to treatment and genotype during cell counting.

### 2.6 Fecal bacterial DNA analysis

#### 2.6.1 Feces collection

Fecal material for qRT-PCR analysis was collected by cutting longitudinally through the colon. Fresh fecal material was removed from the colon then immediately placed into empty microcentrifuge tubes on dry ice. Feces were stored at -80°C until extraction.

#### 2.6.2 Fecal DNA extraction

Fecal DNA extraction was performed using a modified version of the Qiagen PowerLyzer Power Soil Kit as previously reported [41]. Fecal material was added to a PowerBead tube for every animal. DNA was then extracted from the fecal material in line with previous reports. DNA concentration was measured using a NanoDrop, and for analysis purposes, normalized to starting fecal mass.

#### 2.6.3 Quantitative real-time PCR (qRT-PCR) of bacterial phyla

qRT-PCR was completed using phylum-specific and universal bacterial 16S rRNA gene primers (**Table 1**). 10 ng of template DNA was run in triplicate for each sample and primer set. Amplification was monitored in real-time via PowerUp SYBR Green fluorescence signal and the following 3-step protocol: 40 cycles of 10s at 95°C, 10s at 60°C, and 30s at 72°C. The ΔCt method was used to compare across phyla changes between sexes and treatment groups; the ΔΔCt method was used to analyze within phyla changes between sexes and treatment groups.

**Table 1:**
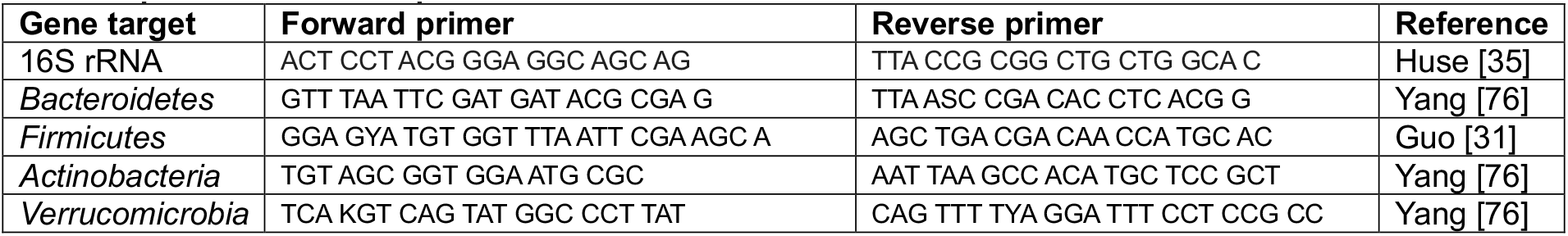
qRT-PCR bacterial primers.

### 2.7 Statistics

All data were analyzed using GraphPad Prism 10. Results were considered statistically significant when *P*<0.05. Results from von Frey tests, dry ice testing, anatomical measures, qPCR, and metabolomics were analyzed with one-, two-, or three-way ANOVA depending on the number of independent variables. If a significant main effect was observed in the ANOVA, Bonferroni post-hoc comparisons were completed.

Microglial staining was analyzed with unpaired t-tests. Needle and paintbrush behaviors were analyzed via Chi-square analyses and Fisher’s post-hoc tests.

## 3. Results

### 3.1 Short-term minocycline administration reduces chronic pain in male, but not female, SCD mice

To determine the analgesic efficacy of minocycline in chronic SCD pain, pain-like behavior testing was completed in male and female SCD mice (HbSS) and hemoglobin control mice (HbAA) after six days of minocycline treatment. Minocycline treatment effectively alleviated punctate mechanical allodynia (**Fig. 1A)**, dynamic mechanical allodynia (**Fig. 1B**), and mechanical hyperalgesia (**Fig. 1C**) in male SCD mice, but had no effect on cold hypersensitivity (**Fig. 1D**). Very different observations were made in female mice. Unlike in male counterparts, minocycline treatment failed to reverse mechanical allodynia (**Fig. 1E, 1F**) or mechanical hyperalgesia (**Fig. 1G**) in female SCD mice; cold hypersensitivity was not observed in this cohort of SCD female mice despite our previous reports clearly demonstrating this phenotype (**Fig. 1H**)[60,61]. Thus, the analgesic efficacy of short-term minocycline treatment in SCD is specific to males.

**Figure 1:**
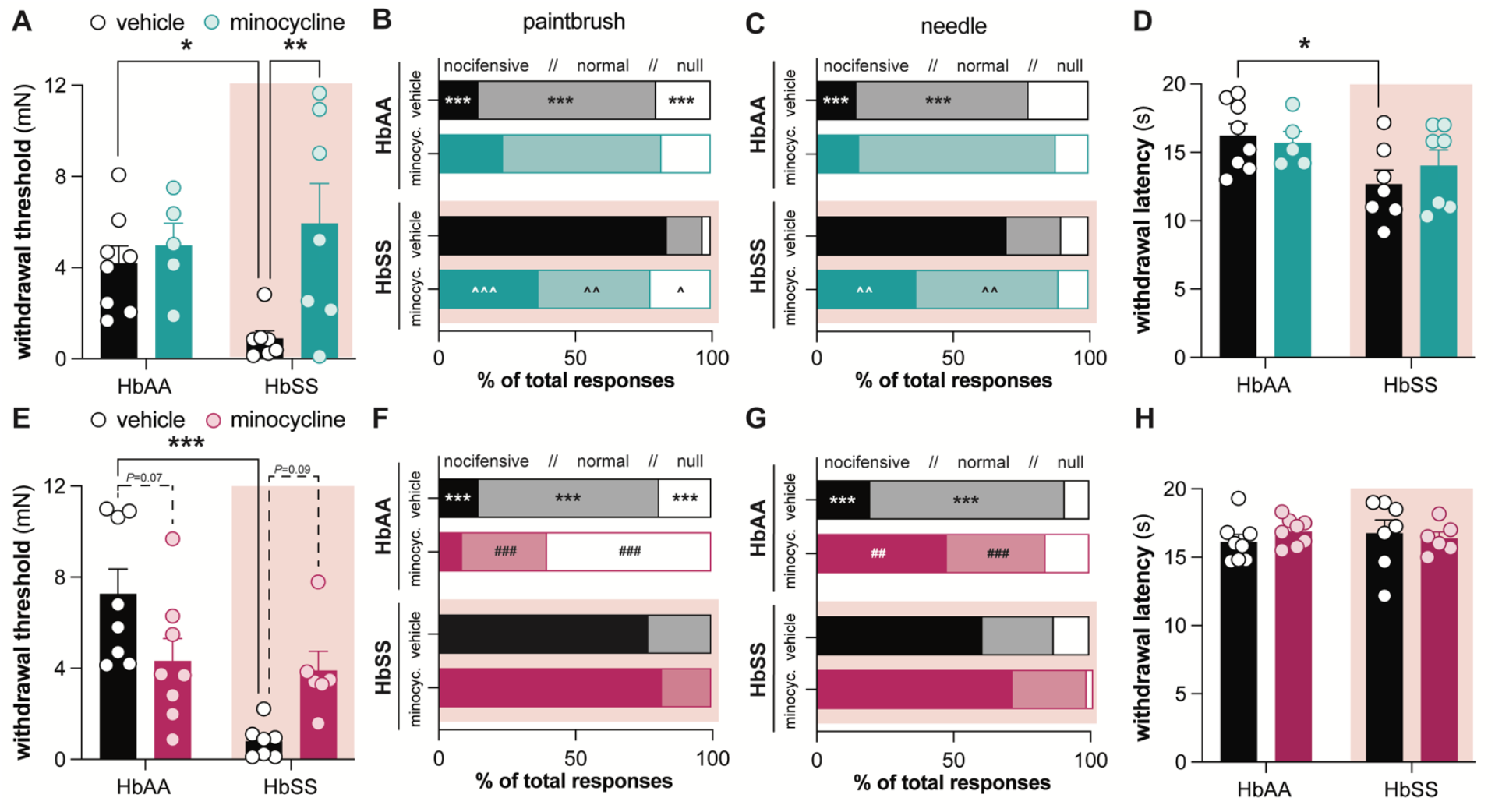
Minocycline alleviates chronic mechanical hypersensitivity in male SCD mice but not in female SCD mice. Male and female SCD mice were maintained on *ad libidum* minocycline treatment (100 mg/kg) for 6 days. Hindpaw **A**. mechanical withdrawal thresholds, **B**. sensitivity to dynamic paintbrush stimulation, **C**. sensitivity to noxious needle stimulation, and **D**. sensitivity to noxious cold stimulation of male mice on day 6 of vehicle or minocycline treatment. Hindpaw **E**. mechanical withdrawal thresholds, **F**. sensitivity to dynamic paintbrush stimulation, **G**. sensitivity to noxious needle stimulation, and **H**. sensitivity to noxious cold stimulation of female mice on day 6 of vehicle or minocycline treatment. *N*=5-8 mice per group; panels B, C, F, G: * vehicle HbAA vs. HbSS, # HbAA vehicle vs. minocycline, ^ HbSS vehicle vs. minocycline.

### 3.2 Minocycline administration has sex-specific effects on pathological organs in SCD mice

We next wanted to determine the manner in which minocycline induced sex-specific analgesia. To begin this analysis, gross anatomical observations were collected for organ systems associated with SCD pathology and pain. Minocycline treatment did not induce significant changes in male mouse body weight (**Fig. 2A**), but it significantly decreased splenomegaly (**Fig. 2B**), a signature of the Townes SCD transgenic mouse model [4]. Increased spleen size in SCD mice may result from accumulation of sickled erythrocytes, bone marrow-independent hematopoiesis, and heightened immune surveillance, particularly of microbial pathogens. Thus, smaller spleen size in minocycline-treated SCD mice may result from a change in gut bacterial populations and resulting immune responses. Minocycline did not affect additional organs of interest in male SCD mice; small intestine and colon length did not differ between genotypes or treatment groups (**Fig. 2C, 2D**).

**Figure 2:**
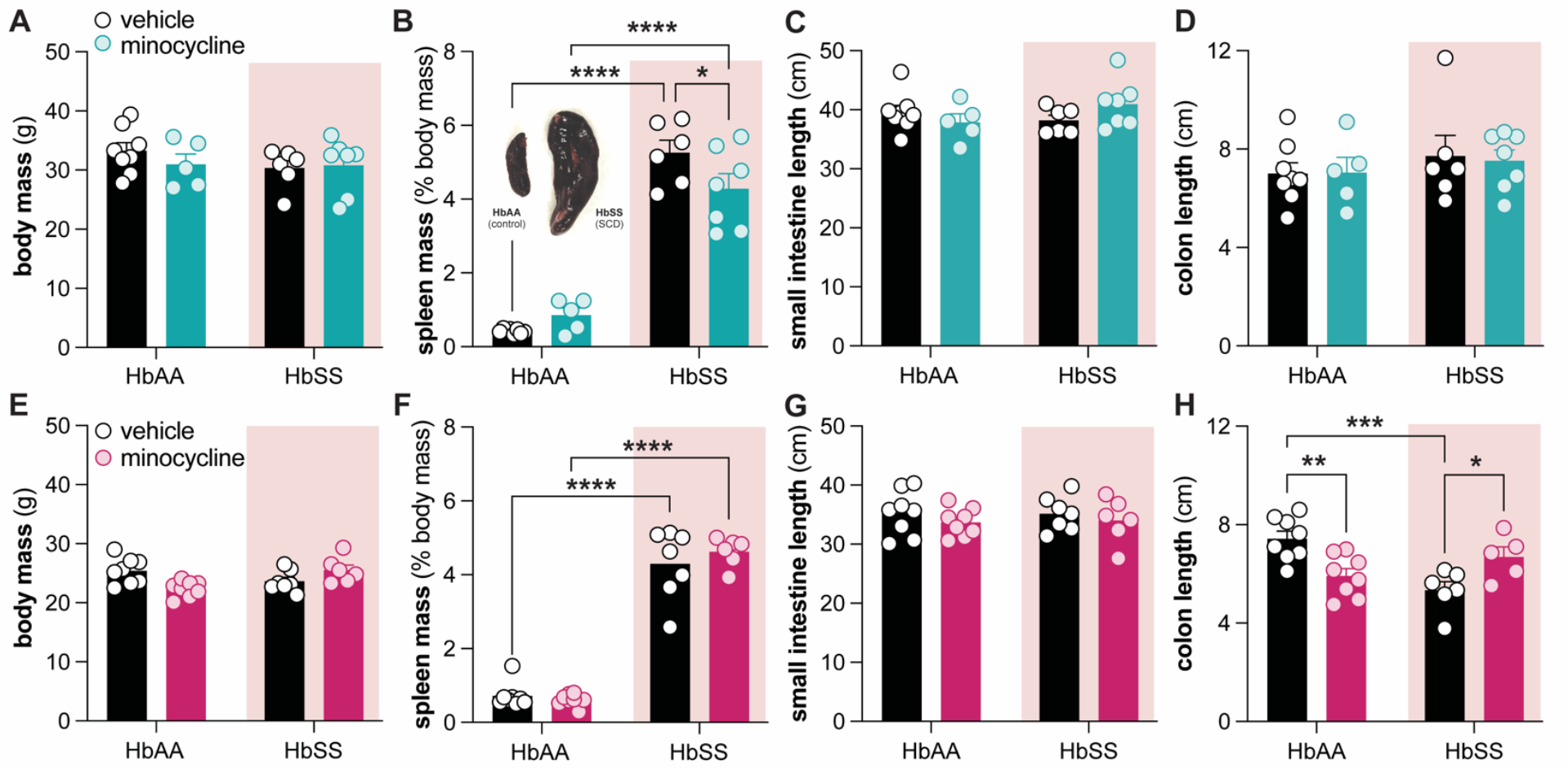
Differential effects of minocycline on gross inflammatory indicators in SCD mice. Male and female SCD (HbSS) and hemoglobin control (HbAA) mice were maintained on *ad libidum* minocycline treatment (100 mg/kg) for 6 days. Gross anatomical observations recorded from male mice on day 6 of treatment included **A**. body mass, **B**. relative spleen mass, **C**. small intestine length, and **D**. colon length. Measures recorded from female mice on day 6 of treatment were identical to those recorded in males and included **E**. body mass, **F**. relative spleen mass, **G**. small intestine length, and **H**. colon length. Bonferroni post-hoc tests: \**P*<0.05, \*\**P*<0.01, \*\*\**P*<0.001, \*\*\*\**P*<0.0001.

The effects of minocycline on female SCD organ systems differed from those observed in males. Minocycline did not change body mass (**Fig. 2E**), relative spleen size (**Fig. 2F**), or small intestine length of female SCD mice (**Fig. 2G**). However, different from male counterparts, female SCD mice had shorter colons than female control mice, a phenotype that was reversed following one week of minocycline treatment (**Fig. 2H**). Colonic shortening is observed in rodent models of intestinal inflammation [27]. Thus, although minocycline treatment does not alleviate reflexive pain measures in female SCD mice, it may still decrease colonic inflammation in a sex-specific manner.

### 3.3 Minocycline analgesia in male SCD mice results, in part, from decreased microglia number in spinal cord

Historically, minocycline analgesia is primarily attributed to its anti-inflammatory properties, most notable of which is its ability to suppress microglia activation in the central nervous system [26]. Intriguingly, functional consequences of this microglial suppression can be sexually dimorphic; despite exhibiting similar levels of injury-induced microgliosis, minocycline treatment induces sex-specific changes in microglial gene expression and metabolite release following injury [2,22,63]. Given this, we hypothesized that the sex-specific analgesic effects of minocycline treatment may result from differential effects on microglial activity in central nociceptive circuits. To assess this possibility, Iba1 immunostaining was performed on lumbar spinal cord isolated from vehicle and minocycline treated SCD mice (**Fig. 3A, 3C, 3E**). More microglia were observed in the dorsal horn of vehicle-treated male SCD mice as compared to vehicle-treated female SCD mice (**Fig. 3B**). Furthermore, minocycline treatment had opposing effects on microglial number in male and female mice; minocycline treatment trended to decrease microglial number in male SCD mice (**Fig. 3D**) while increasing microglial number in female mice (**Fig. 3F**; 2-way ANOVA of data from 3D and 3F combined: overall treatment x sex interaction *P*=0.05). Based on these data, we conclude that SCD factors induce spinal microgliosis in a sex-dependent manner. These effects are further compounded by the sexually dimorphic effects of minocycline treatment on SCD spinal microglia.

**Figure 3:**
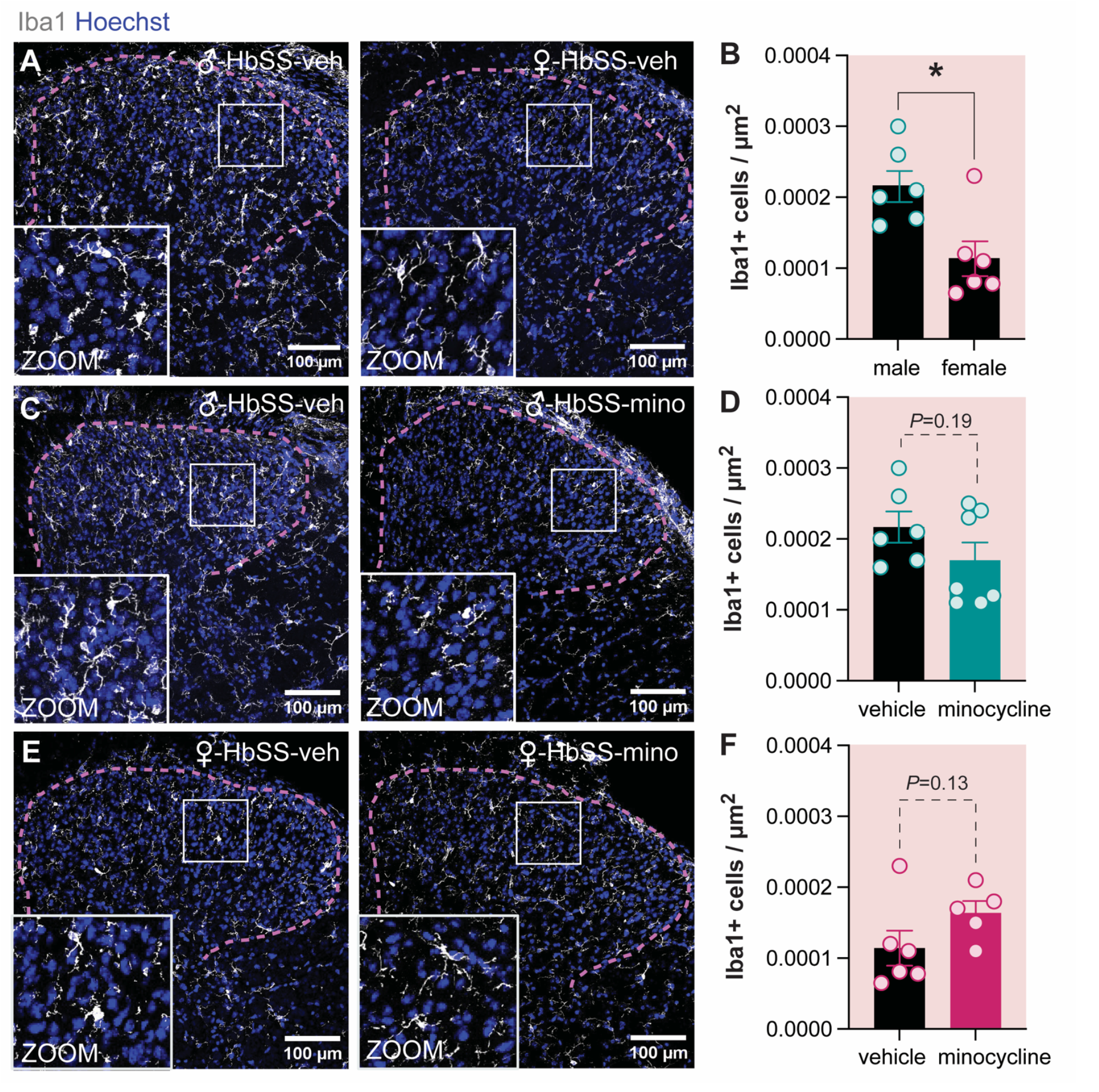
Sex-specific effects of SCD and minocycline treatment on spinal microglia. Representative images (**A, C, E**) and quantification (**B, D, F**) of Iba1+ microglia in the dorsal horn of **A, B**. vehicle-treated male and female HbSS (SCD) mice, **C, D**. vehicle and minocycline-treated male HbSS mice, **E, F**. vehicle and minocycline-treated female HbSS mice. \**P*<0.05, unpaired t-test with Welch’s correction.

### 3.4 Minocycline analgesia results, in part, from effects on male SCD mouse gut microbiome

Given that minocycline had effects on both spleen size and colon length in SCD mice, we reasoned that minocycline-induced changes in the gut microbiome may also contribute to the sex-specific analgesia observed in male SCD mice. To directly test this hypothesis, a series of fecal microbiota transplant (FMT) experiments was performed. In this paradigm, the gut microbiome of sex-matched C57BL/6 mice was altered by oral administration of resuspended fecal material collected from either vehicle- or minocycline-treated SCD mice (**Fig. 4A**). Hindpaw mechanical sensitivity was measured in FMT recipients at various points throughout the paradigm to determine if changes in the gut microbiota alter pain-like behaviors. In line with previous work [39,57], male C57BL/6 mice that received vehicle-treated SCD FMT developed hindpaw mechanical hypersensitivity that persisted for > 1 week following the last FMT. In contrast, male mice that received FMT from minocycline-treated SCD donors did not develop the same mechanical allodynia phenotype (**Fig. 4B, 4C**). Different observations were made in female mice. FMT from both vehicle- and minocycline-treated SCD female mice induced pain in female FMT recipients (**Fig. 4B, 4C**). Notably, mechanical hypersensitivity persisted in female FMT recipients for > 2 weeks, regardless of FMT donor treatment. Thus, in addition to having differential effects on spinal microgliosis, minocycline analgesia in male SCD mice can also be attributed to changes in the gut microbiome that do not occur in female SCD mice.

**Figure 4:**
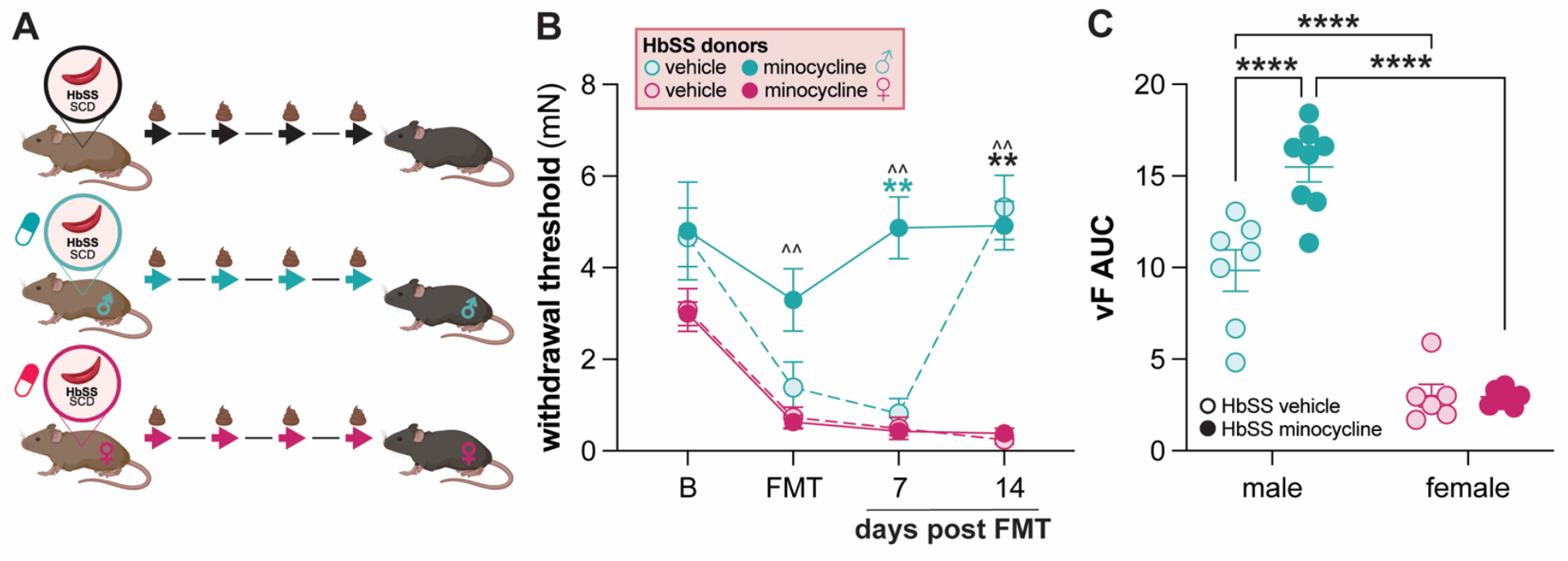
Minocycline-induced changes in the male SCD gut microbiome contribute to sex-specific analgesia. **A**. Schematic of fecal microbiota transplant (FMT) experiments. Fecal material was collected from vehicle- or minocycline-treated SCD (HbSS) mice then orally administered to naïve sex-matched C57BL/6 recipients every other day over the course of 7 days. **B**. Hindpaw mechanical withdrawal thresholds of FMT recipients throughout the paradigm (B: baseline; *N*=6-8; Bonferroni post-hoc tests: minocycline male vs. female ^^*P*<0.01, male minocycline vs. vehicle \*\**P*<0.01, vehicle male vs. female \*\**P*<0.01). **C**. Area under the curve (AUC) calculation for von Frey behavioral assessments during entirety of FMT paradigm.

### 3.5 Minocycline does not exert sex-specific antibiotic effects on bacterial phyla

Given the robust difference in male and female behavior following FMT, we examined whether minocycline treatment induces sex-specific antibiotic effects in SCD mice. Before measuring the relative abundance of individual bacteria, cecum size and fecal DNA content were compared between mice used as FMT donors. Increased cecum size was noted in both minocycline-treated HbAA (hemoglobin control) male (**Fig. 5A**) and female (**Fig. 5B**) mice. This was expected as increased cecum size is a well-documented indicator of antibiotic efficacy. Increased cecum size was not observed in minocycline-treated SCD mice, most likely because vehicle-treated SCD mice trended to have heavier ceca than hemoglobin control mice. In addition to increasing cecum size, minocycline treatment also increased DNA concentrations detected in male hemoglobin control mouse feces, again providing additional support for the antibiotic effects of minocycline (**Fig. 5C**). This same observation was not made in feces collected from female SCD or hemoglobin control mice (**Fig. 5D**).

**Figure 5:**
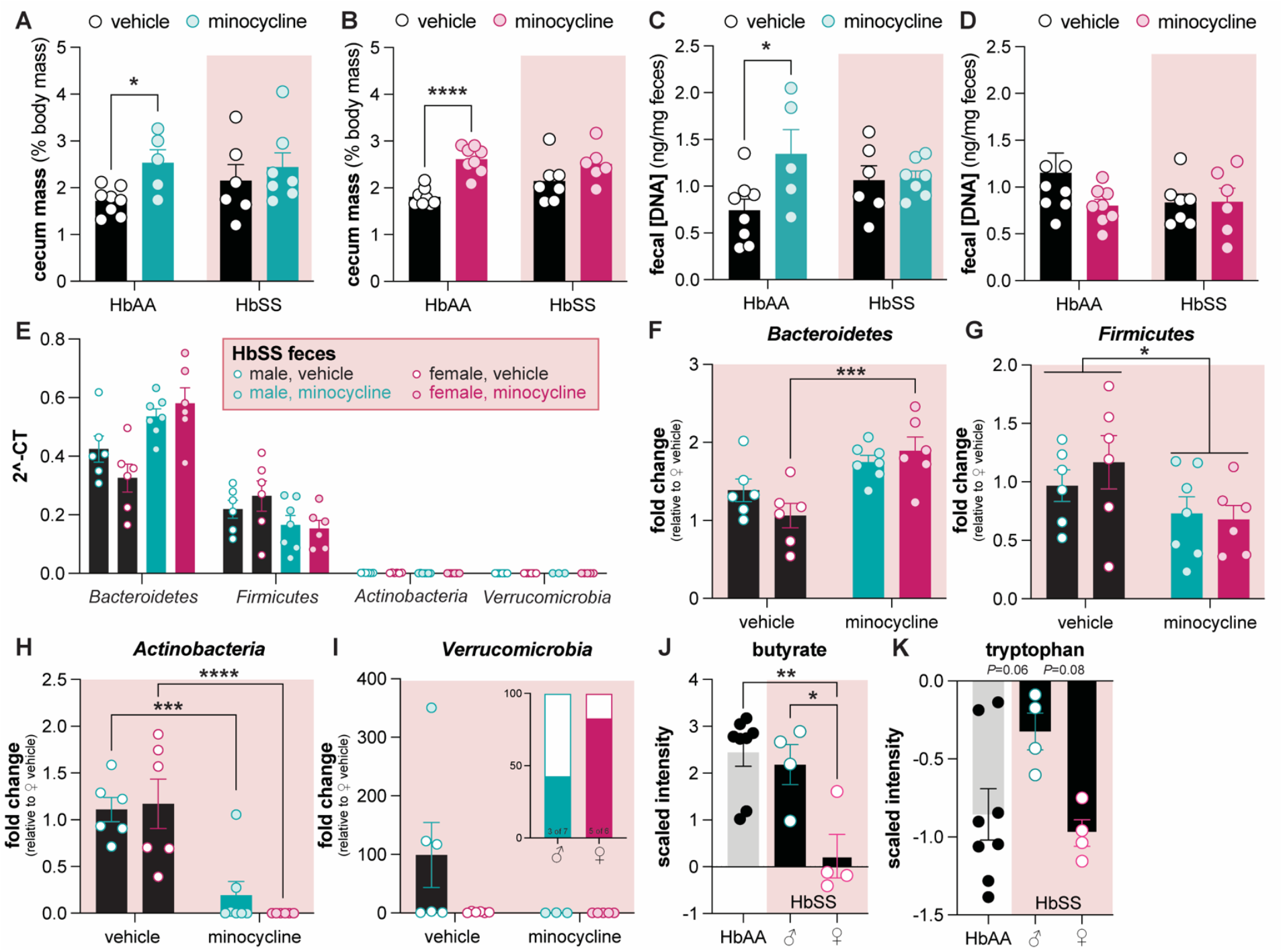
Minocycline-induced changes in the SCD gut microbiota. Male and female SCD (HbSS) and hemoglobin control (HbAA) mice were maintained on *ad libitum* minocycline treatment (100 mg/kg) for 6 days. Following treatment, relative cecum mass for **A**. male and **B**. female mice and DNA content in feces from **C**. male and **D**. female mice was recorded. **E**. Relative abundance of bacterial phyla in feces collected from vehicle and minocycline treated SCD mice. Relative abundance of **F**. *Bacteroidetes*, **G**. *Firmicutes* (2-way ANOVA overall effect of treatment \**P*<0.05), **H**. *Actinobacteria*, and **i**. *Verrucomicrobia* in feces collected from vehicle and minocycline treated SCD mice. Panel J inset illustrates % of minocycline-treated male (teal) and female (magenta) mice with detectable *Verrucomicrobia* in feces. Relative concentration of **J**. butyrate and **K**. tryptophan in feces collected from vehicle-treated HbAA hemoglobin control mice, as well as male and female vehicle-treated HbSS SCD mice. Unless otherwise noted, Bonferroni post-hoc tests: \**P*<0.05, \*\*\**P*<0.001, \*\*\*\**P*<0.0001.

To specifically examine if minocycline alters SCD intestinal bacterial populations in a sex-specific manner, quantitative real-time PCR was used to quantify the relative abundance of the primary bacterial phyla present in mouse intestines. As predicted, *Bacteroidetes* and *Firmicutes* were the most abundant phyla detected in both vehicle- and minocycline-treated SCD feces; members of phyla *Actinobacteria* and *Verrucomicrobia* were present at much lower levels (**Fig. 5E**). A subsequent analysis on individual phyla was completed to better assess if sex-specific effects of minocycline could be detected. Indeed, minocycline treatment significantly increased the abundance of *Bacteroidetes* in female SCD feces but did not have the same effect in male SCD mice (**Fig. 5F**). This was the only sex difference observed in our analysis. Minocycline treatment decreased the relative abundance of *Firmicutes* and *Actinobacteria* in both male and female SCD mice (**Fig. 5G, 5H**). Finally, given the highly variable levels of *Verrucomicrobia* detected across samples, there were no statistical differences noted between treatment groups (**Fig. 5I**). Given the relatively similar effects of minocycline on male and female bacterial populations, we next examined sex-differences in fecal metabolites. In a re-analysis of our previously published data set, we found that 202 of 1030 metabolites detected in SCD mouse feces differed between males and females (**Supp. Table 1**). Of particular note were the short chain fatty acid butyrate and essential amino acid tryptophan. Female SCD mice had significantly less butyrate in their feces as compared to both male SCD mice and HbAA hemoglobin controls (**Fig. 5I**). In contrast, male SCD mice had higher levels of tryptophan in their feces when compared to HbAA controls and female SCD mice (**Fig. 5J**). Future studies should investigate if minocycline treatment affects the abundance of these critical gut metabolites in a sex-specific fashion.

## 4. Discussion

New analgesics are desperately needed for those diagnosed with SCD. To this end, we determined that minocycline may provide chronic pain relief for males suffering from SCD, but not females. This is not the first time male-specific minocycline analgesia has been reported in preclinical pain models. Minocycline effectively alleviates pain in male – but not female – rodents that have received intraplantar injection of formalin [14] or complete Freund’s adjuvant [63], intra-articular injection of HMGB1 [56], collagen antibody-induced arthritis [23], tibia fracture [30], early-life injury-induced priming [51], stress-exacerbated incisional pain [70], chronic constriction injury (CCI) [14,47], and spared nerve injury (SNI) [63].

Although there are many mechanisms through which minocycline can alleviate pain [26], this sex specificity has largely been attributed to the drug’s inhibitory effects on microglia in the male spinal cord. Unlike many previous studies that reported similar levels of microgliosis between sexes following peripheral injury, here we observed sex-specific increases in spinal microgliosis; male SCD mice have more spinal microglia than females. This is the first time this has been reported in transgenic SCD mice. Previous studies have reported increased spinal [66] and hippocampal [32] microgliosis in SCD mice when compared to hemoglobin controls. However, both of these studies only examined tissues from one sex; spinal microgliosis was only assessed in female SCD/control tissue and hippocampal microgliosis was only assessed in male SCD/control tissue. Although not only expressed by microglia, toll-like receptor 4 (TLR4) expression is also increased in SCD mouse spinal cords [43]. In the singular study that reported this finding, both male and female mice were included, but sex-specific analysis of spinal TLR4 expression was not presented [43]. In a similar vein, this study also found that systemic pharmacological inhibition and genetic knockdown of TLR4 alleviated pain in male and female SCD mice, but again, these behavioral results were not segregated by sex. Thus, male-specific increases in spinal microgliosis may be a previously unreported feature in chronic SCD that provides unique opportunities for future therapeutic development.

Although not specifically addressed in our current study, there are several explanations for what could be driving male-specific increases in spinal microgliosis. First is spinal colony-stimulating factor 1 (CSF1) signaling. CSF1 (also known as macrophage colony-stimulating factor; M-CSF) is a cytokine secreted by many cell types, including injured peripheral sensory neurons [29]. When intrathecally injected into naïve mice, CSF1 induces pain and microglial activation, but only in male animals [42]. In female mice, intrathecal CSF1 induces expansion of regulatory T cell (Tregs) populations, cells that subsequently prevent microglial activation [42].

Notably, both CSF1 [45] and Tregs [68] are increased in blood collected from individuals with SCD. CSF1 is also elevated in transgenic SCD mice [45]; to our knowledge, no study has examined Treg populations in SCD animals. Thus, it is possible that in SCD, elevated spinal CSF1 – which may come from either “injured” peripheral nociceptors [59] or additional cell types – selectively induces microglial activation in males.

A second possible mechanism for male-specific microgliosis in SCD is spinal activity of extracellular high mobility group box-1 protein (HMGB1). HMGB1 is classic damage-associated molecular pattern (DAMP). Normally found in the nucleus, HMGB1 is secreted from necrotic cells [62] and released by activated immune cells and peripheral nociceptors [37,73,74]. Notably, individuals and mice with SCD have elevated levels of HMGB1 in circulation that is further increased during acute pain episodes [72]. Previously, intrathecal injection of HMGB1 was found to induce similar pain-like behaviors in male and female mice [1], but *in vitro* exposure to HMGB1 induced sex-specific increases in microglial expression of *Tnf, Ccl2, Il1b*, and *Il6* [2]. Receptors for these factors are expressed by neuronal and immune cells in the dorsal horn – including the microglia from whence they came. Thus, it is possible that in SCD, elevated levels of HMGB1 perpetuate a sex-specific, feed-forward exacerbation of microgliosis that may only be remedied by neutralizing circulating HMGB1.

Although we did not observe a statistically significant decrease in male microglial activity following acute minocycline treatment, it is possible that drug treatment altered the expression and release of compounds from microglia in a sex-dependent manner. Indeed, despite observing similar levels of microgliosis between sexes following peripheral injury, several groups have identified key, sex-specific transcriptional and proteomic changes in the spinal cord following minocycline treatment. For example, in the landmark paper that first described sex-specific immune cell pain modulation, microglial brain-derived neurotrophic factor (BDNF) was critical for pain in male mice but not females [63]. Like every other pro-inflammatory or pro-nociceptive compound mentioned in this manuscript, BDNF is also elevated in SCD plasma [36]. It is unknown if similar increases in microglial-BDNF exist in SCD, but given that the ultimate effect of elevated spinal BDNF signaling is activation of TrkB receptors and subsequent hyperexcitability of dorsal horn neurons – a phenomenon that has been reported in SCD mice [12] – future studies should explore the potential sex-specific analgesic efficacy of TrkB inhibitors in SCD.

In addition to decreased BDNF release in the male spinal cord, minocycline treatment has also been shown to result in male-specific *increases* in spinal haptoglobin and hemopexin [3]. This is incredibly relevant to the current studies given that haptoglobin and hemopexin are, respectively, free hemoglobin and heme scavengers. Individuals and mouse models with SCD have chronically elevated levels of free heme and decreased levels of hemopexin due to the excessive hemolysis that is characteristic of SCD [43,52,67,69].

Elevated heme drives chronic SCD pain by activating TLR4; double transgenic SCD/TLR4 knockout mice are prevented against the development of severe chronic mechanical, thermal, and deep tissue pain and heme-induced exacerbations of this pain [43]. Thus, the male-specific minocycline analgesia observed in the current studies may result from decreased free-heme and subsequent dampening of TLR4-dependent nociceptive signaling in the spinal cord.

Here we demonstrate, for the first time, that sex-specific minocycline analgesia also results from changes in the intestinal milieu. FMT from minocycline-treated male SCD mice did not induce pain in recipients; FMT from vehicle-treated male or female SCD mice as well as minocycline-treated female SCD mice induced mechanical hypersensitivity in recipients, similar to previous reports [39,57]. There are many prokaryotic and eukaryotic factors in the SCD gut that could be impacted by minocycline treatment. First is the gut microbiota, or the bacteria that reside within the intestines. Although we did not observe robust, sex-specific antibiotic effects of minocycline in the current experiments, future studies should more thoroughly test this hypothesis using species-level sequencing. Sex-specific effects of antibiotics have been previously reported in rodents; administration of vancomycin, ciprofloxacin-metronidazole, and a four-drug antibiotic cocktail all lead to sex-specific changes in gut bacteria [25,53] or metabolites [25]. Sex-specific changes in the gut microbiota may also lead to differential effects on the immune system, in particular macrophages which are another known target of minocycline activity [26]. Host and bacterial metabolites are the second factor that may be modulated by minocycline in a sex-dependent fashion. In our previous examination of the SCD mouse gut microbiome, we did not observe sex differences in alpha- or beta-diversity [57]. A secondary analysis of our metabolomic data [57], however, revealed significant differences between relative compound levels in male and female SCD feces. Butyrate and tryptophan are just two of the noteworthy molecules found to be present in different levels in male and female SCD guts. Butyrate is a bacterial-derived short-chain fatty acid that is critical for gut barrier integrity [54]. The critically low levels of butyrate observed in female SCD mice may provide a partial explanation for the shorter colon length observed in these same animals. Given that minocycline reversed this colonic inflammation phenotype, it is possible that levels of butyrate are also increased in feces collected specifically from minocycline-treated female SCD mice. In contrast, male SCD mice had higher levels of tryptophan in their feces when compared to both female SCD mice and hemoglobin controls. Given that tryptophan metabolites have been shown to have antinociceptive and anti-inflammatory properties [46,48,75], this may indicate that insufficient tryptophan metabolism is occurring specifically in the male SCD gut. Minocycline treatment may prevent the growth of tryptophan-metabolizing competitors, thus allowing for increased tryptophan breakdown, and ultimately decreased gut inflammation.

In closing, these studies demonstrate the analgesic efficacy of minocycline in male SCD mice, and, for the first time, imply that the antibiotic effects of minocycline may also lead to sex-specific analgesia. This is perhaps not surprising given the complex multi-directional interactions between the immune system, gut microbiome, and peripheral nervous system (**Fig. 6**). In addition, these results provide critical sex-specific insight into SCD pain biology. The observation that spinal microgliosis specifically occurs in male mice should encourage a re-examination of accepted SCD pain mechanisms on the basis of sex. Uncovering sexually dimorphic pain processes in this disease state will ultimately allow for more effective, personalized analgesics.

**Figure 6:**
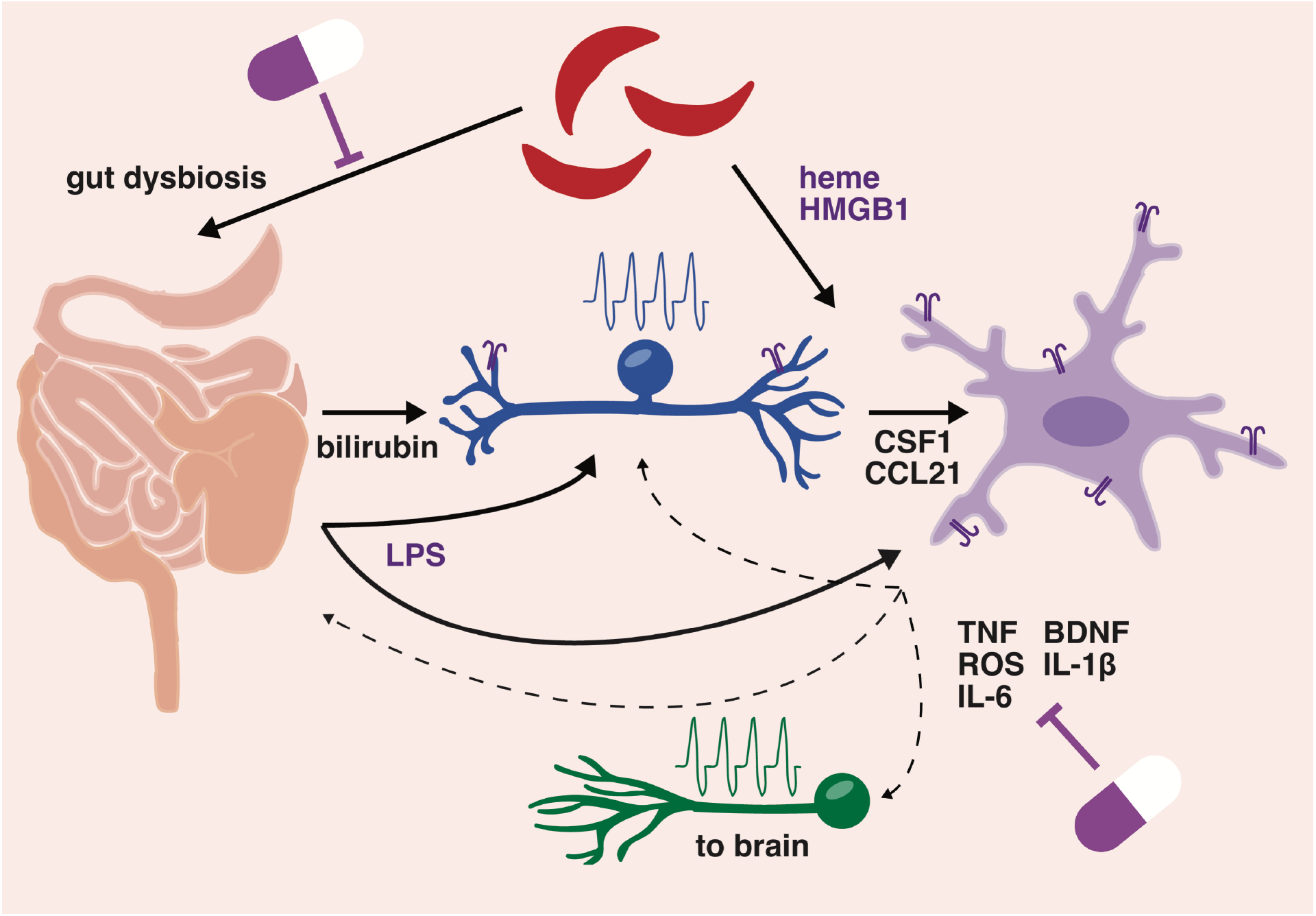
Potential mechanisms of minocycline analgesia in SCD. Summary of how minocycline treatment may influence complex interactions between sickled red blood cells, intestines, primary sensory neurons (blue), spinal microglia (purple), and dorsal horn neurons (green) in SCD. Note that all molecules and features (*i.e*., gut dysbiosis and neuronal activity) listed in image have been reported as elevated in SCD patients or mouse models.

## Acknowledgements

The authors have no conflicts of interest to disclose. The authors would like to thank Nya Gayluak and Victor Cho for assisting with tissue collection, Joe Lombardo and the UT Dallas Imaging and Histology Core for assistance with confocal image collection and analysis, Dr. Michael Burton and lab for Iba1 staining protocol, and Dr. Lena Nguyen for real-time qPCR system and NanoDrop access. Select illustrations were created with BioRender.com. This work was supported by funding from the National Institutes of Health (grant R00 HL155791 to KES) and the Rita Allen Foundation (Award in Pain to KES).

